# DQHTFI: Dynamic-Query Hypergraph Transformer for Fine-Grained Drug–Target Interaction and Affinity Prediction

**DOI:** 10.64898/2026.08.08.743505

**Authors:** Ke Tao, Haoze Chai, Zhuo Chen, Xin Gao, Bin Yu

## Abstract

Drug–target interaction prediction and binding affinity prediction are two key tasks in drug discovery and drug repurposing. Although deep learning methods have made significant progress, existing models typically rely on global representations of drugs and proteins, making it difficult to adequately model fine-grained interactions between their local units. Fixed multimodal fusion strategies also struggle to dynamically adjust the contributions of different modalities for different drug–target combinations. To address these issues, we propose DQHTFI, a fine-grained interaction prediction framework for drug–target interaction classification and binding affinity regression. DQHTFI employs BRICS fragments and Pfam functional domains as the basic interaction units and jointly learns semantic and structural representations. We design a dynamic-query hypergraph Transformer framework in which hyperedges are constructed among the multimodal features of fragment–domain pairs. Dynamic queries are generated from the cross-conditioned features of fragment–domain pairs to adaptively adjust the contribution of each modality, thereby modeling higher-order interactions between local units. Our proposed model achieves competitive results on multiple benchmark datasets.

## Introduction

Drug–target interactions underlie therapeutic efficacy, drug selectivity, and potential adverse effects. They are also central to understanding disease mechanisms, identifying therapeutic targets, and advancing drug repurposing(Yang et al. 2026). In drug discovery, drug–target interaction (DTI) prediction aims to determine whether a candidate drug interacts with a target, whereas drug–target affinity (DTA) prediction further quantifies the binding strength between them. Therefore, accurately modeling drug–target interactions and binding affinities is important for drug development, drug repurposing, and precision medicine.

Traditional studies of drug–target relationships generally rely on in vitro experiments, high-throughput screening, and affinity measurements. Although these approaches can provide reliable experimental evidence, they are often costly, time-consuming, and limited in throughput, making it difficult to efficiently screen large numbers of drug–target combinations. To improve screening efficiency, molecular docking, similarity-based inference, and machine learning methods based on handcrafted features have been widely applied to drug–target prediction. However, their performance often depends on prior interaction knowledge or the quality of manually designed features (Ye et al. 2021). With the increasing availability of drug–target data and advances in deep learning, data-driven approaches have gradually become an important direction for modeling drug–target relationships. Compared with traditional methods, deep learning models can automatically learn latent representations from diverse data modalities, including SMILES sequences, molecular graphs, molecular fingerprints, amino acid sequences, and structural information, while capturing complex nonlinear relationships between drugs and targets through end-to-end training.

Despite the substantial progress of deep learning methods in DTI and DTA prediction, several limitations remain. For DTI prediction, some methods incorporate multimodal information, such as sequence and graph-structured representations of drugs and targets, but typically rely on feature concatenation or global fusion strategies (Bian et al. 2024; Zhai et al. 2025), limiting their ability to exploit modality complementarity and adapt fusion to different drug–target combinations (Yang et al. 2026). For DTA prediction, many methods mainly use sequence-level or global representations, making it difficult to capture fine-grained associations between local drug and target units and affinity differences arising from different combinations of local structures (He et al. 2025a; Lv et al. 2025). From a biological perspective, the activity of a small-molecule drug is closely related to the physicochemical properties and structural characteristics of specific chemical fragments, whereas protein recognition and function frequently depend on domains, binding pockets, or other local functional regions (Kumar, Romano, and Ritchie 2025; Lv et al. 2025). Therefore, modeling fine-grained associations between local drug structures and functional target regions can more comprehensively characterize drug–target relationships and improve model interpretability.

To address these issues, we propose DQHTFI, a fine-grained drug–target prediction framework for DTI classification and DTA regression. For drug representation, DQHTFI decomposes molecules into local chemical fragments using BRICS and employs ChemBERTa and GINE to learn the semantic and graph-structural representations of these fragments, respectively. For target representation, DQHTFI identifies functional protein domains using Pfam, obtains domain-level semantic representations with ESM-2, and employs a graph attention mechanism to learn associations among domains. Based on these representations, we construct hyperedges at the fragment–domain level and design a dynamic-query hypergraph Transformer. Dynamic queries are generated from the cross-conditioned features of fragment–domain pairs to adaptively adjust the contribution of each modality, thereby modeling higher-order interactions among local units. A low-rank bilinear interaction mechanism is subsequently used to identify and aggregate key local interactions for DTI classification and DTA regression.

The main contributions are three-fold as follows:

- We propose DQHTFI, which uses BRICS drug fragments and Pfam protein domains as basic interaction units and explicitly models the potential associations between fragments and domains.
- We design a dynamic-query hypergraph Transformer module. Hyperedges are constructed among the multiple modalities of each fragment–domain pair, and dynamic queries are generated based on the cross-conditioned features to adaptively fuse multimodal features.
- We design a bilinear local interaction aggregation module that scores and performs weighted aggregation over fragment–domain pairs to form a drug–target interaction representation.

## Related Work

### Multimodal Representation Learning

Early studies on drug–target relationship prediction mainly relied on traditional machine learning methods and hand-crafted features. Drugs were commonly represented using molecular fingerprints, chemical descriptors, or similarity profiles, whereas proteins were encoded using amino acid compositions or sequence-derived statistical features. Their predictive performance therefore largely depended on feature quality, making it difficult to fully capture the complex nonlinear relationships between drug structures and protein sequences (Chen et al. 2016; Bagherian et al. 2021).

With the development of deep learning, research gradually shifted toward end-to-end representation learning. DeepConv-DTI employs convolutional neural networks to extract local residue patterns from protein sequences (Lee, Keum, and Nam 2019). DeepDTA directly learns continuous representations from drug SMILES and protein sequences (Öztürk, Özgür, and Ozkirimli 2018), whereas GraphDTA represents drugs as molecular graphs to capture molecular topology (Nguyen et al. 2021). More recently, pretrained models such as ChemBERTa and ESM-2 have enhanced the representation of chemical language and contextual information in protein sequences (Tang, Zhao, and Wang 2025; Wu et al. 2026). These advances enable models to integrate sequence, graph-structural, and pretrained features. However, existing multimodal methods often encode different modalities independently before downstream fusion, leaving the complementary relationships among modalities insufficiently exploited (Yang et al. 2026).

### Drug–Target Interaction Fusion Modeling

After obtaining drug and target representations, modeling their potential interactions is a key step in both DTI classification and DTA regression (Wang et al. 2025b; Duan et al. 2026). Early deep learning methods mainly used feature concatenation, vector products, or multilayer perceptrons to combine drug and target representations. Although these strategies can capture overall associations, they provide limited characterization of complex interactions between local drug and protein features (Gao and Zhu 2026; Hua et al. 2025b).

To improve interaction modeling, HyperAttentionDTI employs multidimensional interaction attention to model fine-grained associations between atoms and amino acids (Zhao et al. 2022). DrugBAN adopts a bilinear attention network to capture pairwise feature associations between drugs and targets (Bai et al. 2023), while DACMF-DTI introduces a dual-attention framework for cross-modal fusion (Wei et al. 2025). DrugKANs and KAN-MoDTI further employ Kolmogorov– Arnold networks to enhance nonlinear interaction representation (Fu et al. 2025; Liu et al. 2026). These studies demonstrate the effectiveness of attention mechanisms and bilinear interactions in modeling drug–target matching relationships. Despite these advances, many methods still rely on fixed or global fusion strategies and cannot adaptively adjust the contributions of different modalities for specific drug– target combinations (Yang et al. 2026). Some affinity prediction methods also incorporate binding pockets or three-dimensional structural information to enhance interaction modeling. However, their applicability may be limited when reliable structural data are unavailable or contain substantial prediction errors (He et al. 2025b; Lv et al. 2025).

### Fine-Grained Interaction Modeling

Drug–target interaction signals are often concentrated in local drug substructures and functional protein regions rather than global representations alone. The biological activity of a drug is frequently associated with particular chemical fragments, functional groups, or molecular motifs, whereas protein recognition and function commonly depend on domains, binding pockets, or key residue regions (Noble, Endicott, and Johnson 2004). Therefore, local interaction modeling has become an important direction for characterizing drug–target relationships.

Existing studies have explored fine-grained modeling from different perspectives. MolTrans extracts local patterns from drug and protein sequences through frequent subsequence mining and models their pairwise relationships (Huang et al. 2021). SP-DTI incorporates protein subpocket information to characterize local spatial regions in targets (Liu et al. 2025). MotifGT-DTI uses molecular motifs to enhance drug graph representations (Tian et al. 2026a), while MSN-DTA extracts multiscale structural information from drug molecules and employs attention to identify important substructures (Wang et al. 2025a). These methods improve the representation of local structural information beyond sequence-level or molecule-level features.

However, existing fine-grained approaches either depend on binding pockets or three-dimensional structural priors,or focus primarily on drug substructures and residue-level interactions (Zhu et al. 2025; Yang et al. 2026). Consequently, explicit pairwise interactions between drug fragments and protein functional domains remain insufficiently explored.

## Method

### Model Overview

DQHTFI is a fine-grained drug–target prediction framework for DTI classification and DTA regression. As illustrated in Figure 1, drug molecules are decomposed into BRICS fragments, while functional domains are identified from protein sequences using Pfam. Semantic and graph-structural representations are then extracted for both fragments and domains. Based on these representations, a dynamic-query hypergraph Transformer adaptively models multimodal interactions within each fragment–domain pair. Bilinear local interaction aggregation subsequently identifies and aggregates important local interactions to obtain the final drug–target representation.The pseudocode for DQHTFI is provided in the appendix.

**Figure 1.**
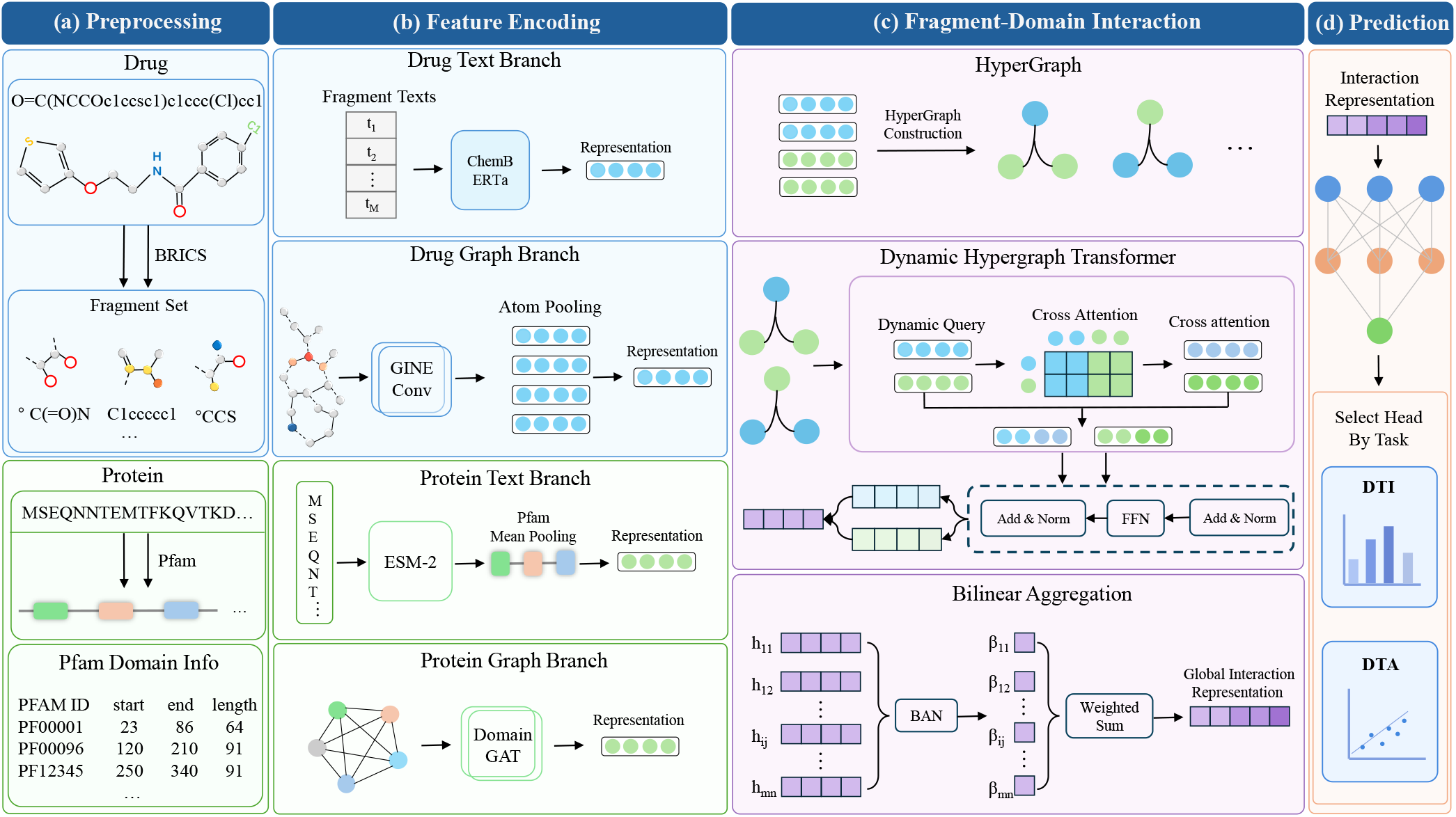
Overall framework of DQHTFI, which comprises four main stages: (a) An input and preprocessing module, which processes drug SMILES and protein sequences into BRICS fragments, molecular graphs, and Pfam domains. (b) A feature encoding module, which extracts semantic and graph-structural representations of fragments and domains. (c) A fragment– domain interaction module, which models local interactions through hypergraph construction, a dynamic-query hypergraph Transformer, and bilinear aggregation. (d) A prediction module for DTI classification and DTA regression.

### Fine-Grained Feature Encoding

#### Drug fragment encoding

Given the SMILES representation *S* of a drug molecule, BRICS (Degen et al. 2008) decomposes it into the fragment set F (*S*) = {*f*_1_, *f*_2_, …, *f*_*M*_}. Here, *M* denotes the number of fragments. ChemBERTa (Chithrananda, Grand, and Ramsundar 2020) encodes each fragment SMILES. Its token representations are aggregated through attention-mask-aware mean pooling and then projected to obtain the semantic representation 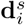 ∈ *R*^*d*^ of the *i*-th fragment.

To capture the structural context of each fragment within the complete molecule, the drug is further represented as a molecular graph *G* = (*V, E*). GINEConv (Hu et al. 2020) performs message passing using atom and bond features. The resulting atom representations are mean-pooled according to the atom-to-fragment mapping obtained from BRICS decomposition, yielding the fragment graph representation 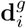∈ *R*^*d*^.

#### Protein domain encoding

Given a target protein sequence *P* = {*a*_1_, *a*_2_, …, *a*_*L*_}, Pfam (Mistry et al. 2021) identifies the domain set D (*P*) = *δ*_1_, {*δ*_1,_*δ*_2_, …, *δ*_*N*_}. Here, *N* denotes the number of domains. ESM-2 (Lin et al. 2023) encodes the complete sequence to obtain contextual residue representations. For each domain, the representations between its start and end positions are mean-pooled to obtain a domain-level embedding. An independent projection then produces the domain semantic representation 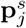 ∈*R*^*d*^.

For the graph branch, the same domain-level ESM-2 embeddings are independently projected as initial node features **x**_*j*_ ∈*R*^*d*^. We construct a fully connected directed domain graph with self-loops. For an edge from domain *a* to domain *b*, the relation descriptor **r**_*ab*_ contains the source and destination features, their absolute difference, and their element-wise product. A two-layer MLP produces the head-specificrelation bias 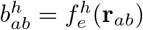

Let 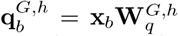 and 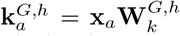 The edge aware attention coefficient is calculated as

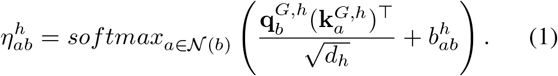

The neighborhood (*b*) contains all domain nodes, including node *b* itself. Multi-head aggregation followed by residual normalization produces the domain graph representation 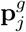 ∈ *R*^*d*^.

### Dynamic-Query Hypergraph Transformer

Each drug fragment is paired with each protein domain to form a local hyperedge containing four modality-specific representations: 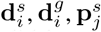, and 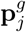. The complete hyperedge set contains all valid fragment–domain combinations. Instead of assigning the same fixed queries to all hyperedges, DQHTFI generates pair-specific queries from their multimodal features.

For each fragment–domain pair, four cross-conditioned features are constructed through element-wise multiplication as

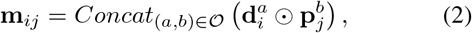

where *O* = (*s, s*), (*s, g*), (*g, s*), (*g, g*) specifies the concatenation order, ⊙ denotes element-wise multiplication, and **m**_*ij*_ ∈*R*^4*d*^.

The four original representations and **m**_*ij*_ are concatenated in the order 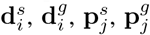 and **m**_*ij*_. This produces the pair-conditioned representation **c**_*ij*_ ∈ *R*^8*d*^. Two independent two-layer MLPs then generate the dynamic queries as

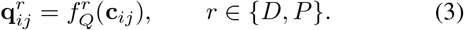

The indices *D* and *P* denote the drug-side and protein-side branches, respectively. The query networks 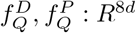→ *R*^*d*^ produce one *d*-dimensional query for each branch.

The four modality-specific representations are stacked as row tokens and combined with learnable modality-role embeddings:

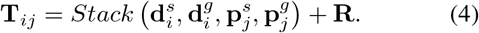

Thus, **T**_*ij*_ ∈ *R*^4×*d*^. Its rows correspond to the drug semantic, drug structural, protein semantic, and protein structural modalities. The matrix **R** ∈ *R*^4×*d*^ contains the corresponding learnable role embeddings.

For the *h*-th attention head, define 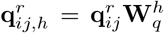 and 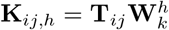 The attention weights are calculated as

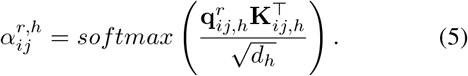

The softmax operation is performed over the four modality tokens. The drug-side and protein-side queries independently attend to the same token set.

Let 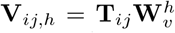 The corresponding multi-head attention context is

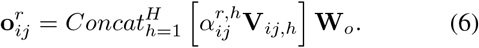

Dropout is applied to the attention weights and attention contexts during training.

Each dynamic query and its attention context are subsequently processed by a standard post-norm Transformer block. A residual attention update is followed by a residual feed-forward network, with layer normalization applied after each residual connection. The resulting drug-side and protein-side contexts are denoted by 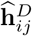 and 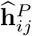, respectively.

This operation is performed independently within each fragment–domain hyperedge. The contributions of different hyperedges are coordinated by the subsequent bilinear aggregation module.

The ba<u>s</u>e fragment and domain representations are calculated as 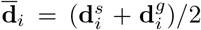 and 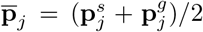. For compact notation, let 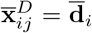 and 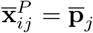 The dynamic contexts are incorporated through residual fusion:

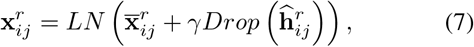

where *r* ∈ {*D, P*}, *LN* and *Drop* denote layer normalization and dropout, respectively, and *γ* is a learnable scaling parameter. For subsequent interaction modeling, we use 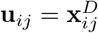 and 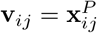

The local interaction operator *ϕ*(**u, v**) concatenates the two input representations, their element-wise product, and their absolute difference. The local interaction representation is

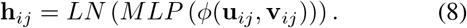

Specifically, *ϕ*(**u**_*ij*_, **v**_*ij*_) contains **u**_*ij*_, **v**_*ij*_, **u**_*ij*_ ⊙ **v**_*ij*_, and |**u**_*ij*_ − **v**_*ij*_ |. The representation **h**_*ij*_ characterizes the fine-grained interaction between the *i*-th drug fragment and the *j*-th protein domain.

### Bilinear Local Interaction Aggregation

Different fragment–domain pairs may contribute differently to the final drug–target prediction. DQHTFI therefore employs a low-rank bilinear attention mechanism (Bai et al. 2023) to score and aggregate the local interaction representations.

The pair-conditioned representations are projected into a low-rank interaction space as ũ_*ij*_ = *Drop*(**u**_*ij*_**U**) and 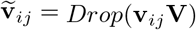 The projection matrices satisfy **U, V**∈ *R*^*d*×*ρ*^, where *ρ* denotes the low-rank projection dimension.

The interaction score is calculated as

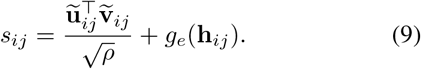

The bilinear term captures the multiplicative association between the pair-conditioned fragment and domain representations. The two-layer MLP *g*_*e*_() independently evaluates the encoded local interaction.

The scores are normalized over all valid fragment–domain pairs:

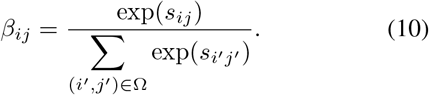

The set Ω contains all valid fragment–domain pairs after padded positions are excluded. Therefore, ∑ _(*i,j*)_ Ω *β*_*ij*_ = 1.

The final drug–target interaction representation is obtained through weighted aggregation:

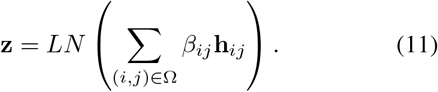

The normalized weight *β*_*ij*_ represents the contribution of the corresponding local interaction, and **z** is the global interaction representation used for prediction.

### Prediction and Optimization

Since DTI and DTA have different task objectives, separate prediction heads are constructed on the shared interaction representation **z**.

#### DTI prediction

DTI prediction is formulated as a binary classification task with *y*∈ {0, 1}. The DTI prediction head produces the interaction logit *l* = *MLP*_DTI_(**z**). The model is optimized using binary cross-entropy with logits. During evaluation and inference, the corresponding interaction probability is obtained as 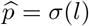.

#### DTA prediction

DTA prediction is formulated as a regression task with *y* ∈ *R*. The DTA prediction head produces the continuous affinity value *ŷ* = *MLP*_DTA_(**z**). The model is optimized using mean squared error.

## Experiments

### Experimental Setup

#### Datasets

For DTI prediction, we conduct experiments on four public benchmark datasets: BindingDB (Liu et al. 2007), BioSNAP (Zitnik et al. 2018), and the Human and C.elegans datasets (Liu et al. 2015). Under the regular setting, each dataset is randomly divided into training, validation, and test sets at a ratio of 8:1:1. We further adopt a dual cold-start setting to evaluate model generalization. AUROC, AUPR, and F1 are used as evaluation metrics, and the model with the highest validation AUROC is selected for testing.

For DTA prediction, we conduct experiments on the Davis (Davis et al. 2011) and KIBA (Tang et al. 2014) datasets. Under the regular setting, each dataset is randomly divided into training, validation, and test sets at a ratio of 8:1:1. We further consider drug cold-start and target cold-start settings. CI, MSE, and 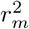 are used to evaluate affinity prediction performance, and the model with the highest validation CI is selected for testing.

#### Baselines

To comprehensively evaluate the predictive performance of DQHTFI on DTI classification and DTA regression, we select a range of representative deep learning methods as baselines. For DTI prediction, the compared methods include GraphDTA (Nguyen et al. 2021), HyperAttDTI (Zhao et al. 2022), MolTrans (Huang et al. 2021), DrugBAN (Bai et al. 2023), MotifGT-DTI (Tian et al. 2026a), and MMDG-DTI (Hua et al. 2025a); for DTA prediction, the compared methods include GraphDTA (Nguyen et al. 2021), DeepDTA (Öztürk, Özgür, and Ozkirimli 2018), MGraphDTA (Yang et al. 2022), MFCLDTA (Tian et al. 2026b), DeepDTAGen (Shah et al. 2025), and MT-DiffGen (Li et al. 2026). These methods cover diverse technical paradigms, including sequence modeling, molecular graph learning, attention mechanisms, Transformers, multifeature and multimodal fusion, contrastive learning, and generative modeling. For all baseline methods, we preferentially use their publicly available official implementations and reproduce them using the hyperparameter settings recommended in the original papers or the default configurations provided in the official code.Detailed descriptions of the baseline methods are provided in the appendix.

#### Implementation Details

Our model is implemented using PyTorch, with PyTorch Geometric employed to construct the molecular graph and protein domain graph encoding modules. All experiments are conducted on a Linux server equipped with an NVIDIA GeForce RTX 3090 GPU. The model is trained using the AdamW optimizer. Early stopping is applied based on validation performance. All experiments are independently repeated five times, and the average results over the five runs are reported.Detailed experimental settings and hyperparameter configurations are provided in the appendix.

### Performance Evaluation

#### DTI Regular Experiments

Table 1 presents the results on four DTI datasets under the regular setting. DQHTFI achieves the best AUROC, AUPR, and F1 scores on both BindingDB and BioSNAP. Specifically, it obtains scores of 0.962, 0.958, and 0.898 on BindingDB, and 0.928, 0.930, and 0.862 on BioSNAP, respectively. Compared with the best baselines, DQHTFI improves these metrics by up to 1.1%, 0.4%, and 0.9%. On C.elegans and Human, DQHTFI achieves the best F1 and AUROC, respectively, while remaining close to the best results on the other metrics. Overall, DQHTFI demonstrates stable and competitive performance across datasets with different scales and distributions.

**Table 1:** Performance comparison of DQHTFI and the baseline methods on four DTI datasets under the regular setting. **Bold** and <u>underlined</u> values denote the best and second-best results, respectively.

| Methods | BindingDB |  |  | BioSNAP |  |  | C.elegans |  |  | Human |  |  |
| --- | --- | --- | --- | --- | --- | --- | --- | --- | --- | --- | --- | --- |
|  | AUROC | AUPR | F1 | AUROC | AUPR | F1 | AUROC | AUPR | F1 | AUROC | AUPR | F1 |
| GraphDTA | 0.945 | 0.935 | 0.862 | 0.883 | 0.886 | 0.785 | 0.972 | 0.872 | 0.917 | 0.953 | 0.954 | 0.892 |
| HyperAttDTI | 0.957 | 0.945 | 0.882 | 0.900 | 0.898 | 0.834 | 0.987 | 0.987 | <u>0.955</u> | 0.984 | 0.981 | 0.942 |
| MolTrans | 0.948 | 0.936 | 0.861 | 0.890 | 0.896 | 0.820 | 0.980 | 0.982 | <u>0.952</u> | 0.974 | 0.972 | 0.941 |
| DrugBAN | 0.956 | 0.945 | 0.881 | 0.902 | 0.901 | 0.830 | 0.983 | 0.984 | 0.947 | 0.981 | 0.980 | 0.937 |
| MotifGT-DTI | 0.956 | 0.952 | 0.890 | 0.918 | 0.926 | 0.852 | 0.991 | 0.989 | 0.950 | 0.990 | **0.989** | 0.948 |
| MMDG-DTI | 0.960 | 0.954 | 0.889 | 0.872 | 0.865 | 0.859 | **0.994** | **0.993** | 0.953 | 0.986 | 0.987 | **0.950** |
| DQHTFI | **0.962** | **0.958** | **0.898** | **0.928** | **0.930** | **0.862** | 0.990 | 0.988 | **0.956** | **0.991** | 0.986 | 0.946 |

#### DTI Cold-Start Experiments

Table 2 presents the results on BindingDB and BioSNAP under the dual cold-start setting. On BindingDB, DQHTFI achieves AUROC and AUPR scores of 0.730 and 0.649, improving the best baselines by 2.1% and 0.9%, respectively. Its F1 score reaches 0.594 and remains competitive with the best-performing method. On BioSNAP, DQHTFI obtains the best AUROC, AUPR, and F1 scores of 0.848, 0.847, and 0.764, corresponding to improvements of 1.8%, 3.2%, and 2.0%. These results indicate that DQHTFI maintains strong classification performance even when the training and test sets contain no overlapping drugs or targets. This demonstrates its generalization ability for previously unseen drug–target combinations.

**Table 2:** Performance comparison under the DTI dual cold-start setting. **Bold** and <u>underlined</u> values denote the best and second-best results, respectively.

| Methods | BindingDB |  |  | BioSNAP |  |  |
| --- | --- | --- | --- | --- | --- | --- |
|  | AUROC | AUPR | F1 | AUROC | AUPR | F1 |
| GraphDTA | 0.584 | 0.580 | 0.526 | 0.656 | 0.662 | 0.579 |
| HyperAttDTI | 0.659 | 0.596 | 0.580 | 0.731 | 0.757 | 0.625 |
| MolTrans | 0.593 | 0.550 | 0.509 | 0.669 | 0.695 | 0.610 |
| DrugBAN | 0.653 | 0.614 | 0.541 | 0.649 | 0.664 | 0.633 |
| MotifGT-DTI | 0.682 | 0.631 | 0.590 | 0.758 | 0.752 | 0.738 |
| MMDG-DTI | 0.715 | 0.643 | **0.602** | 0.833 | 0.821 | 0.749 |
| DQHTFI | **0.730** | **0.649** | <u>0.594</u> | **0.848** | **0.847** | **0.764** |

#### DTA Regular Experiments

Table 3 compares DQHTFI with the baseline methods on Davis and KIBA under the regular setting. On Davis, DQHTFI obtains CI, MSE, and 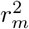 values of 0.898, 0.213, and 0.696, respectively. Its CI is tied for the best result, while its MSE is the lowest among all compared methods, demonstrating strong ranking performance and low prediction error. On KIBA, DQHTFI achieves the second-best CI of 0.874, while its MSE and 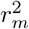values are 0.172 and 0.658. Overall, DQHTFI demonstrates strong performance on Davis and competitive ranking performance on KIBA, although its KIBA regression accuracy can be further improved.

**Table 3:** Performance comparison on Davis and KIBA under the regular DTA setting. **Bold** and <u>underlined</u> values denote the best and second-best results, respectively.

| Methods | Davis |  |  | KIBA |  |  |
| --- | --- | --- | --- | --- | --- | --- |
| | CI | MSE | $r_m^2$ | CI | MSE | $r_m^2$ |
| GraphDTA | 0.868 | 0.264 | 0.635 | 0.846 | 0.244 | 0.588 |
| DeepDTA | 0.872 | 0.251 | 0.630 | 0.853 | 0.231 | 0.650 |
| MGraphDTA | 0.871 | 0.227 | 0.705 | 0.855 | 0.173 | 0.763 |
| MFCLDTA | <u>0.892</u> | 0.220 | **0.732** | **0.882** | <u>0.168</u> | **0.793** |
| DeepDTAGen | **0.898** | 0.215 | 0.693 | 0.871 | 0.169 | 0.680 |
| MT-DiffGen | 0.887 | <u>0.215</u> | 0.705 | 0.868 | **0.160** | 0.702 |
| DQHTFI | **0.898** | **0.213** | 0.696 | <u>0.874</u> | 0.172 | 0.658 |

#### DTA Cold-Start Experiments

Tables 4 and 5 present the results under the drug cold-start and target cold-start settings, respectively. Under drug cold-start, DQHTFI achieves the best performance across all three metrics on Davis, with CI, MSE, and 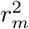 values of 0.767, 0.595, and 0.357, respectively. On KIBA, it obtains the highest CI of 0.756 and the lowest MSE of 0.382, although its 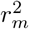 does not reach the best result. Under target cold-start, DQHTFI achieves the highest CI and 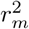on Davis, reaching 0.826 and 0.487, respectively, while its MSE ranks second. On KIBA, DQHTFI also obtains the highest CI of 0.683, but its MSE and 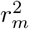 remain below the best-performing results. Overall, DQHTFI demonstrates strong and stable ranking ability under both cold-start settings. Its advantages are particularly evident on Davis and in the CI metric, while the regression accuracy on KIBA still has room for improvement.

**Table 4:** Performance comparison on Davis and KIBA under the drug cold-start setting. **Bold** and <u>underlined</u> values denote the best and second-best results, respectively.

| Methods | Davis |  |  | KIBA |  |  |
| --- | --- | --- | --- | --- | --- | --- |
| | CI | MSE | $r_m^2$ | CI | MSE | $r_m^2$ |
| GraphDTA | 0.665 | 0.802 | 0.146 | 0.670 | 0.585 | 0.211 |
| DeepDTA | 0.688 | 0.773 | 0.158 | 0.686 | 0.548 | 0.241 |
| MGraphDTA | 0.704 | 0.708 | 0.179 | 0.707 | 0.449 | 0.350 |
| MFCLDTA | 0.693 | 0.623 | 0.203 | <u>0.748</u> | 0.435 | 0.391 |
| DeepDTAGen | 0.744 | 0.680 | 0.251 | 0.741 | 0.420 | **0.440** |
| MT-DiffGen | 0.732 | 0.616 | <u>0.258</u> | 0.735 | 0.468 | 0.393 |
| DQHTFI | **0.767** | **0.595** | **0.357** | **0.756** | **0.382** | 0.284 |

**Table 5:** Performance comparison on Davis and KIBA under the target cold-start setting. **Bold** and <u>underlined</u> values denote the best and second-best results, respectively.

| Methods | Davis |  |  | KIBA |  |  |
| --- | --- | --- | --- | --- | --- | --- |
| | CI | MSE | $r_m^2$ | CI | MSE | $r_m^2$ |
| GraphDTA | 0.684 | 0.776 | 0.241 | 0.597 | 0.652 | 0.193 |
| DeepDTA | 0.702 | 0.760 | 0.278 | 0.611 | 0.619 | 0.200 |
| MGraphDTA | 0.803 | 0.585 | 0.396 | 0.632 | 0.588 | 0.359 |
| MFCLDTA | 0.817 | **0.429** | <u>0.440</u> | <u>0.679</u> | <u>0.461</u> | **0.406** |
| DeepDTAGen | 0.811 | 0.640 | 0.355 | 0.674 | **0.433** | 0.396 |
| MT-DiffGen | 0.820 | 0.657 | 0.339 | 0.664 | 0.485 | <u>0.381</u> |
| DQHTFI | **0.826** | <u>0.565</u> | **0.487** | **0.683** | 0.495 | 0.196 |

### Ablation Study

To evaluate the contribution of each component in DQHTFI, we conduct DTI classification and DTA regression ablation experiments on the BindingDB and Davis datasets, respectively. As illustrated in Figure 2, the complete model is denoted as S0, while each remaining variant removes or replaces one key component. The dataset partitioning, training strategy, and other parameter settings are kept unchanged across all variants.

**Figure 2.**
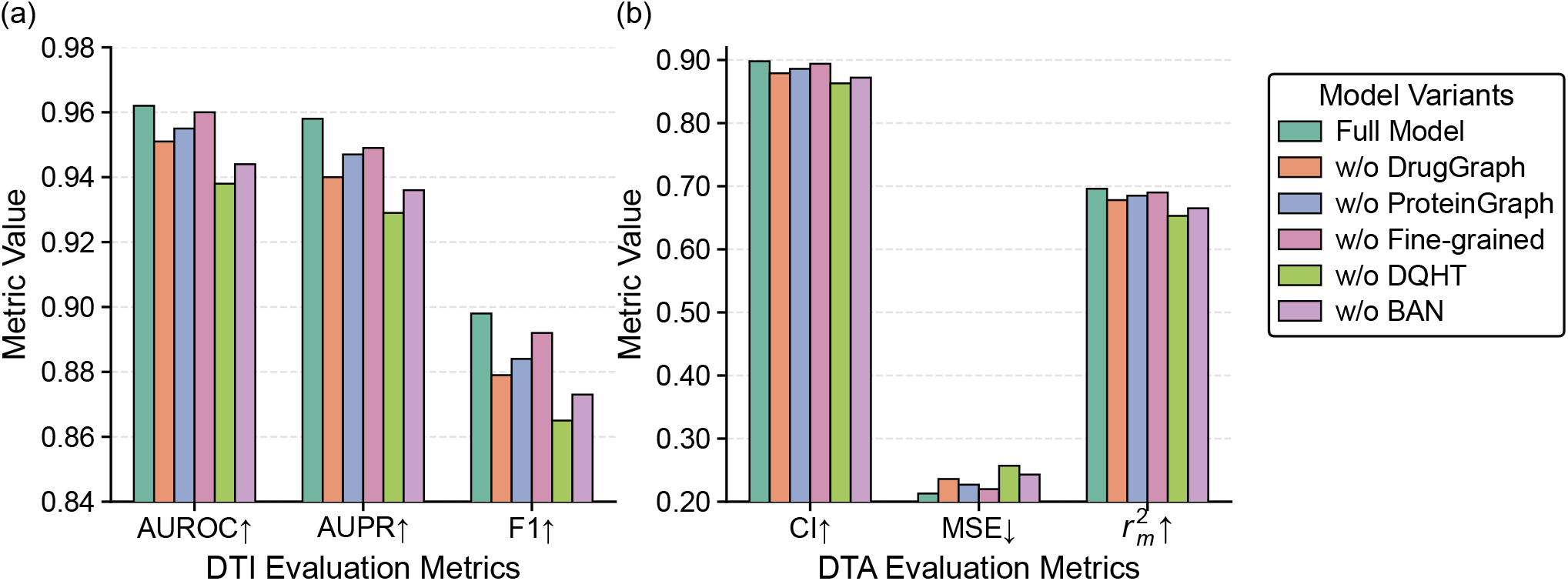
Ablation study results of DQHTFI. (a) DTI classification on BindingDB. (b) DTA regression on Davis.

#### S1: w/o Drug Graph

We remove the drug graph branch and retain only the semantic representations of drug fragments. Compared with the complete model, AUROC, AUPR, and F1 decrease by 1.1%, 1.9%, and 2.1% on the DTI task, respectively. On the DTA task, CI and 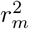 decrease by 2.1% and 2.6%, respectively, while MSE increases by 10.8%. These results indicate that the drug graph branch provides effective molecular structural information complementary to the fragment representations.

#### S2: w/o Protein Graph

We remove the protein domain graph branch and retain only the semantic representations of protein domains. AUROC, AUPR, and F1 decrease by 0.7%, 1.1%, and 1.6% on the DTI task, respectively. On the DTA task, CI and 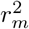 decrease by 1.3% and 1.6%, respectively, while MSE increases by 6.6%. These results demonstrate that associations among protein domains provide complementary information to their local semantic representations.

#### S3: w/o Fine-Grained Interaction

We separately aggregate the drug fragment and protein domain representations and replace the fine-grained fragment–domain interactions with a global drug–target interaction. AUROC, AUPR, and F1 decrease by 0.2%, 0.9%, and 0.7% on the DTI task, respectively. On the DTA task, CI and 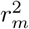 decrease by 0.4% and 0.9%, respectively, while MSE increases by 3.3%. These results indicate that fine-grained interaction modeling maintains the predictive performance of the model while supporting interpretability analysis at the fragment–domain level.

#### S4: w/o DQHT

We replace the dynamic-query hypergraph Transformer with a multilayer perceptron. This variant produces the most significant performance degradation. AU-ROC, AUPR, and F1 decrease by 2.5%, 3.0%, and 3.7% on the DTI task, respectively. On the DTA task, CI and 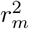 decrease by 3.9% and 6.2%, respectively, while MSE increases by 20.7%. These results demonstrate that DQHT effectively models conditional multimodal associations across different fragment–domain combinations.

#### S5: w/o BAN

We replace the bilinear interaction aggregation module with mean pooling. AUROC, AUPR, and F1 decrease by 1.9%, 2.3%, and 2.8% on the DTI task, respectively. On the DTA task, CI and 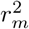decrease by 2.9% and 4.5%, respectively, while MSE increases by 14.1%. These results indicate that weighted interaction aggregation more effectively distinguishes the contributions of different local interactions to the final prediction.

## Conclusion

In this work, we propose DQHTFI, a fine-grained framework for both DTI classification and DTA regression. DQHTFI uses BRICS drug fragments and Pfam protein domains as local interaction units. It learns semantic and graph-structural representations for both fragments and domains. For each fragment–domain pair, the corresponding multimodal features are organized into a hyperedge. A dynamic-query hypergraph Transformer then adaptively fuses these features and captures higher-order local associations. A low-rank bilinear aggregation module further scores and combines the local interactions into a global representation. Experimental results on multiple DTI and DTA benchmark datasets show that DQHTFI achieves competitive performance under both regular and cold-start settings. Ablation studies further confirm the contributions of the graph-structural branches, the dynamic-query hypergraph Transformer, and the bilinear interaction aggregation module.

## Notes

### Competing Interest Statement

The authors have declared no competing interest.

## References

Bagherian, M.; Sabeti, E.; Wang, K.; Sartor, M. A.; Nikolovska-Coleska, Z.; and Najarian, K. 2021. Machine Learning Approaches and Databases for Prediction of Drug– Target Interaction: A Survey Paper. Briefings in Bioinformatics, 22: 247–269.

Bai, P.; Miljković, F.; John, B.; and Lu, H. 2023. Interpretable Bilinear Attention Network with Domain Adaptation Improves Drug–Target Prediction. Nature Machine Intelligence, 5: 126–136.

Bian, J.; Lu, H.; Dong, G.; and Wang, G. 2024. Hierarchical Multimodal Self-Attention-Based Graph Neural Network for DTI Prediction. Briefings in Bioinformatics, 25: bbae293.

Chen, X.; Yan, C. C.; Zhang, X.; Zhang, X.; Dai, F.; Yin, J.; and Zhang, Y. 2016. Drug–Target Interaction Prediction: Databases, Web Servers and Computational Models. Briefings in Bioinformatics, 17: 696–712.

Chithrananda, S.; Grand, G.; and Ramsundar, B. 2020. ChemBERTa: Large-Scale Self-Supervised Pretraining for Molecular Property Prediction. arXiv:2010.09885.

Davis, M. I.; Hunt, J. P.; Herrgard, S.; Ciceri, P.; Wodicka, L. M.; Pallares, G.; Hocker, M.; Treiber, D. K.; and Zarrinkar, P. P. 2011. Comprehensive Analysis of Kinase Inhibitor Selectivity. Nature Biotechnology, 29(11): 1046–1051.

Degen, J.; Wegscheid-Gerlach, C.; Zaliani, A.; and Rarey, M. 2008. On the Art of Compiling and Using “Drug-Like” Chemical Fragment Spaces. ChemMedChem, 3: 1503–1507.

Duan, H.; Dong, J.; Zhao, Y.; Wang, S.; and Wang, W. 2026. Multi-View Fusion Feature Representation Learning for Drug-Target Interaction Prediction. Knowledge-Based Systems, 337: 115364.

Fu, X.; Du, Z.; Chen, Y.; Chen, H.; Zhuo, L.; Lu, A.; Cao, D.; and Yao, X. 2025. DrugKANs: A Paradigm to Enhance Drug-Target Interaction Prediction with KANs. IEEE Journal of Biomedical and Health Informatics, 1–12.

Gao, D.; and Zhu, F. 2026. HMT-DTI: Hierarchical Meta-Path Learning with Transformer for Drug–Target Interaction Prediction. Neural Networks, 194: 108093.

He, C.; Yang, C.; Zhang, H.; Long, Y.; and Zhao, X. 2025a. DTI-MPFM: A Multi-Perspective Fusion Model for Predicting Potential Drug–Target Interactions. Expert Systems with Applications, 264: 125740.

He, H.; Chen, G.; Tang, Z.; and Chen, C. Y.-C. 2025b. Dual Modality Feature Fused Neural Network Integrating Binding Site Information for Drug Target Affinity Prediction. npj Digital Medicine, 8: 67.

Hu, W.; Liu, B.; Gomes, J.; Zitnik, M.; Liang, P.; Pande, V.; and Leskovec, J. 2020. Strategies for Pre-Training Graph Neural Networks. In International Conference on Learning Representations.

Hua, Y.; Feng, Z.; Song, X.; Wu, X.-J.; and Kittler, J. 2025a. MMDG-DTI: Drug–Target Interaction Prediction via Multi-modal Feature Fusion and Domain Generalization. Pattern Recognition, 157: 110887.

Hua, Y.; Xu, T.; Song, X.; Feng, Z.; Wang, R.; Zhang, W.; and Wu, X. 2025b. R-DTI: Drug Target Interaction Prediction Based on Second-Order Relevance Exploration. In Proceedings of the AAAI Conference on Artificial Intelligence, volume 39, 17368–17376.

Huang, K.; Xiao, C.; Glass, L. M.; and Sun, J. 2021. MolTrans: Molecular Interaction Transformer for Drug– Target Interaction Prediction. Bioinformatics, 37: 830–836.

Kumar, R.; Romano, J. D.; and Ritchie, M. D. 2025. CASTER-DTA: Equivariant Graph Neural Networks for Predicting Drug–Target Affinity. Briefings in Bioinformatics, 26(5): bbaf554.

Lee, I.; Keum, J.; and Nam, H. 2019. DeepConv-DTI: Prediction of Drug-Target Interactions via Deep Learning with Convolution on Protein Sequences. PLOS Computational Biology, 15: e1007129.

Li, S.; Wang, H.; Hu, T.; Zhuang, L.; Zhao, J.; Sheng, Y.; and Sun, Y. 2026. MT-DiffGen: Unifying Affinity Prediction and Target-Aware Molecule Generation with a Multi-Task Diffusion Model. Knowledge-Based Systems, 340: 115605.

Lin, Z.; Akin, H.; Rao, R.; Hie, B.; Zhu, Z.; Lu, W.; Smetanin, N.; Verkuil, R.; Kabeli, O.; Shmueli, Y.; dos Santos Costa, A.; Fazel-Zarandi, M.; Sercu, T.; Candido, S.; and Rives, A. 2023. Evolutionary-Scale Prediction of Atomic-Level Protein Structure with a Language Model. Science, 379(6637): 1123–1130.

Liu, H.; Jia, H.; Li, W.; Li, W.; and Yuan, Y. 2026. KAN-MoDTI: Drug Target Interaction Prediction Based on Kolmogorov-Arnold Network and Multimodal Feature Fusion. Expert Systems with Applications, 298: 129828.

Liu, H.; Sun, J.; Guan, J.; Zheng, J.; and Zhou, S. 2015. Improving Compound–Protein Interaction Prediction by Building Up Highly Credible Negative Samples. Bioinformatics, 31(12): i221–i229.

Liu, S.; Liu, Y.; Xu, H.; Xia, J.; and Li, S. Z. 2025. SP-DTI: Subpocket-Informed Transformer for Drug–Target Interaction Prediction. Bioinformatics, 41: btaf011.

Liu, T.; Lin, Y.; Wen, X.; Jorissen, R. N.; and Gilson, M. K. 2007. BindingDB: A Web-Accessible Database of Experimentally Determined Protein–Ligand Binding Affinities. Nucleic Acids Research, 35(suppl_1): D198–D201.

Lv, T.; Zhu, J.; Liu, J.; Nie, S.; Tian, H.; Xiao, Y.; Liu, Y.; Li, L.; and Pan, X. 2025. M 2N: A Progressive Macro-to-Micro 3D Modeling Scheme for Unveiling Drug–Target Affinity. In Proceedings of the AAAI Conference on Artificial Intelligence, volume 39, 586–594.

Mistry, J.; Chuguransky, S.; Williams, L.; Qureshi, M.; Salazar, G. A.; Sonnhammer, E. L. L.; Tosatto, S. C. E.; Paladin, L.; Raj, S.; Richardson, L. J.; Finn, R. D.; and Bateman, A. 2021. Pfam: The Protein Families Database in 2021. Nucleic Acids Research, 49: D412–D419.

Nguyen, T.; Le, H.; Quinn, T. P.; Nguyen, T.; Le, T. D.; and Venkatesh, S. 2021. GraphDTA: Predicting Drug–Target Binding Affinity with Graph Neural Networks. Bioinformatics, 37(8): 1140–1147.

Noble, M. E. M.; Endicott, J. A.; and Johnson, L. N. 2004. Protein Kinase Inhibitors: Insights into Drug Design from Structure. Science, 303: 1800–1805.

Öztürk, H.; Özgür, A.; and Ozkirimli, E. 2018. DeepDTA: Deep Drug–Target Binding Affinity Prediction. Bioinformatics, 34: i821–i829.

Shah, P. M.; Zhu, H.; Lu, Z.; Wang, K.; Tang, J.; and Li, M. 2025. DeepDTAGen: A Multitask Deep Learning Framework for Drug-Target Affinity Prediction and Target-Aware Drugs Generation. Nature Communications, 16: 5021.

Tang, J.; Szwajda, A.; Shakyawar, S.; Xu, T.; Hintsanen, P.; Wennerberg, K.; and Aittokallio, T. 2014. Making Sense of Large-Scale Kinase Inhibitor Bioactivity Data Sets: A Comparative and Integrative Analysis. Journal of Chemical Information and Modeling, 54(3): 735–743.

Tang, W.; Zhao, Q.; and Wang, J. 2025. LLMDTA: Improving Cold-Start Prediction in Drug–Target Affinity with Biological LLM. IEEE Transactions on Computational Biology and Bioinformatics, 22(6): 2398–2409.

Tian, W.; Zeng, M.; Wang, J.; and Lu, C. 2026a. MotifGT-DTI: Pivotal Motif-Based Graph Transformer Model Improves Drug–Target Interaction Prediction. IEEE Transactions on Neural Networks and Learning Systems, 1–15.

Tian, Z.; Zhu, S.; Teng, Z.; Yan, X.; and Wang, T. 2026b. MFCLDTA: Multi-Scale Feature Contrastive Learning for Predicting Drug-Target Binding Affinity. Expert Systems with Applications, 306: 130918.

Wang, J.; Ding, P.; Zhu, Y.; Gao, X.; Yu, X.; and Yu, B. 2025a. MSN-DTA: A Multi-Scale Node Adaptive Graph Neural Network for Interpretable Drug-Target Binding Affinity Prediction. Knowledge-Based Systems, 320: 113699.

Wang, M.; Lei, X.; Guo, L.; Chen, M.; and Pan, Y. 2025b. DHGT-DTI: Advancing Drug-Target Interaction Prediction through a Dual-View Heterogeneous Network with Graph-SAGE and Graph Transformer. Journal of Pharmaceutical Analysis, 15: 101336.

Wei, Z.; Zheng, X.; Li, C.; Wang, M.; and Tang, C. 2025. DACMF-DTI: Dual Attention Embedded Cross-Modality Fusion for Drug-Target Interaction Prediction. Knowledge-Based Systems, 326: 114063.

Wu, Q.; Lv, J.; Zhang, Z.; and Cui, F. 2026. Generalizable Drug–Target Interaction Prediction via ESM-2 Representations and Progressive Contrastive Curriculum Learning. In Proceedings of the AAAI Conference on Artificial Intelligence, volume 40, 1276–1284.

Yang, J.; Zhang, J.; Qian, K.; Yang, Q.; Li, W.; and Cheng, Z. 2026. Bridging the Modality Reliability Gap in Drug–Target Interaction Prediction via a Confidence-Aware Multimodal Fusion Framework. In Proceedings of the AAAI Conference on Artificial Intelligence, volume 40, 27529–27537.

Yang, Z.; Zhong, W.; Zhao, L.; and Chen, C. Y.-C. 2022. MGraphDTA: Deep Multiscale Graph Neural Network for Explainable Drug–Target Binding Affinity Prediction. Chemical Science, 13: 816–833.

Ye, Q.; Hsieh, C.-Y.; Yang, Z.; Kang, Y.; Chen, J.; Cao, D.; He, S.; and Hou, T. 2021. A Unified Drug–Target Interaction Prediction Framework Based on Knowledge Graph and Recommendation System. Nature Communications, 12: 6775.

Zhai, X.; Wang, C.; Wang, R.; Kang, J.; Li, S.; Chen, B.; Ma, T.; Zhou, Z.; Yang, C.; and Shi, C. 2025. Blend the Separated: Mixture of Synergistic Experts for Data-Scarcity Drug– Target Interaction Prediction. In Proceedings of the AAAI Conference on Artificial Intelligence, volume 39, 22336–22344.

Zhao, Q.; Zhao, H.; Zheng, K.; and Wang, J. 2022. HyperAttentionDTI: Improving Drug–Protein Interaction Prediction by Sequence-Based Deep Learning with Attention Mechanism. Bioinformatics, 38: 655–662.

Zhu, Z.; Ding, Y.; Qi, G.; Cong, B.; Li, Y.; Bai, L.; and Gao, X. 2025. Drug–Target Affinity Prediction Using Rotary Encoding and Information Retention Mechanisms. Engineering Applications of Artificial Intelligence, 147: 110239.

Zitnik, M.; Sosič, R.; Maheshwari, S.; and Leskovec, J. 2018. BioSNAP Datasets: Stanford Biomedical Network Dataset Collection. Stanford Biomedical Network Dataset Collection.

