## Supplementary Appendix for "DQHTFI: Dynamic-Query Hypergraph Transformer for Fine-Grained Drug–Target Interaction and Affinity Prediction"

### A Training Procedure

Algorithm 1 summarizes the end-to-end training procedure of DQHTFI, including feature encoding, dynamic-query fragment–domain interaction modeling, bilinear aggregation, and task-specific prediction. Binary cross-entropy with logits is used for DTI classification, whereas mean squared error is used for DTA regression.

**Algorithm 1** DQHTFI.

**Require:** Preprocessed splits $\mathcal{D}_\tau^{\mathrm{tr}}$, $\mathcal{D}_\tau^{\mathrm{val}}$, and $\mathcal{D}_\tau^{\mathrm{te}}$, task type $\tau$, and epochs $E$

Initialize model parameters $\theta$
**for** $epoch = 1$ to $E$ **do**
  **for** each batch $(\mathcal{B}, Y)$ in $\mathcal{D}_\tau^{\mathrm{tr}}$ **do**
    $X_D = (H_D^s, H_D^g) \leftarrow \mathrm{DrugEncode}(\mathcal{B})$
    $X_P = (H_P^s, H_P^g) \leftarrow \mathrm{TargetEncode}(\mathcal{B})$
    $m_{ij} \leftarrow \mathrm{Cat}_{(a,b)\in\mathcal{O}}(d_i^a \odot p_j^b)$
    $c_{ij} \leftarrow \mathrm{Cat}(d_i^s, d_i^g, p_j^s, p_j^g, m_{ij})$
    $Q_{ij} \leftarrow (q_{ij}^D, q_{ij}^P), \quad q_{ij}^r = f_Q^r(c_{ij})$
    $U, V \leftarrow \mathrm{DQHT}(X_D, X_P, \{Q_{ij}\}, \Omega)$
    $h_{ij} \leftarrow \mathrm{PairEncode}(u_{ij}, v_{ij})$
    $s_{ij} \leftarrow \dfrac{\langle W_u u_{ij}, W_v v_{ij}\rangle}{\sqrt{\rho}} + g_e(h_{ij})$
    $\beta_{ij} \leftarrow \dfrac{\exp(s_{ij})}{\sum_{(i',j')\in\Omega}\exp(s_{i'j'})}$
    $z \leftarrow \mathrm{LN}\left(\sum_{(i,j)\in\Omega} \beta_{ij} h_{ij}\right)$
    $\hat{Y}_\tau, L \leftarrow \mathrm{TaskSpecificPredict}(z, Y, \tau)$
    $\theta \leftarrow \mathrm{AdamWUpdate}(\theta, \nabla_\theta L)$
  **end for**
**end for**
$\theta^* \leftarrow \mathrm{SelectBest}(\{\theta_e\}_{e=1}^E, \mathcal{D}_\tau^{\mathrm{val}}, \tau)$
$\hat{Y}_\tau \leftarrow \mathrm{Infer}(\theta^*, \mathcal{D}_\tau^{\mathrm{te}})$
**return** $\hat{Y}_\tau$

### B Dataset Descriptions

For the DTI task, we employ four public benchmark datasets, namely BindingDB, BioSNAP, C. elegans, and Human. BindingDB (Liu et al. 2007) is a large-scale database that collects experimentally measured protein–ligand binding data. The version used in this study contains 14,643 drugs, 2,622 proteins, and 49,113 drug–target pairs. BioSNAP (Zitnik et al. 2018) integrates drug–target association information from STITCH and DrugBank and contains 4,502 drugs, 2,181 proteins, and 27,464 pairs. The C. elegans dataset (Liu et al. 2015) focuses on the model organism *Caenorhabditis elegans* and contains 1,767 drugs, 1,876 proteins, and 7,786 pairs. The Human dataset (Liu et al. 2015) integrates information from DrugBank, Matador, and STITCH and is constructed through confidence-score filtering and negative-sample dissimilarity rules. It contains 2,726 drugs, 2,001 proteins, and 6,728 pairs. To reduce the influence of class imbalance on model training and evaluation, the numbers of positive and negative samples are kept approximately equal in all four DTI datasets.

For the DTA task, we employ the widely used Davis (Davis et al. 2011) and KIBA (Tang et al. 2014) affinity prediction datasets. Davis contains 68 drugs, 442 targets, and 30,056 affinity measurements. Its original labels are dissociation constants $K_d$ measured in nanomolar units. We convert them into logarithmic affinity values according to $\mathrm{p}K_d = -\log_{10}(K_d/10^9)$, resulting in a value range of approximately 5.0–10.8. KIBA contains 260 drugs, 193 targets, and 30,110 drug–target pairs. Its labels are KIBA scores that integrate multiple kinase-inhibitor bioactivity measurements, including $K_i$, $K_d$, and $\mathrm{IC}_{50}$. The score values range approximately from 0.0 to 17.2.

### C Experimental Settings

The experiments were conducted under Ubuntu 20.04 on a server equipped with an NVIDIA GeForce RTX 3090 GPU with 24 GB of GPU memory and 32 GB of system memory. The maximum number of training epochs was set to 200 for the regular DTI and DTA experiments and 100 for all cold-start experiments. The principal hyperparameters are summarized in Table 1.

Table 1: Principal hyperparameter settings of DQHTFI.

| Hyperparameter | Value | Description |
|---|---|---|
| Embedding dimension | 128 | Common feature dimension |
| Learning rate | $1 \times 10^{-4}$ | Initial learning rate |
| Batch size | 128 | Number of samples per batch |
| Dropout | 0.1 | Dropout rate |
| Weight decay | 0.01 | Weight-decay coefficient |
| GINE layers | 2 | Number of molecular graph layers |
| Domain graph layers | 2 | Number of protein domain graph layers |
| Domain graph heads | 4 | Number of domain graph attention heads |
| Bilinear rank | 32 | Low-rank interaction dimension |

### D Baseline Descriptions

This section provides detailed descriptions of the baseline methods used in the experiments.

**GraphDTA** (Nguyen et al. 2021). GraphDTA is a pioneering graph-based framework that represents drug molecules as molecular graphs. It employs graph neural networks to capture the spatial and topological information of drugs, while retaining a CNN branch for protein sequence encoding.

**HyperAttDTI** (Zhao et al. 2022). HyperAttDTI employs a multidimensional interaction attention mechanism to model fine-grained associations between drug atoms and protein residues, thereby enhancing the representation of potential drug–protein interactions.

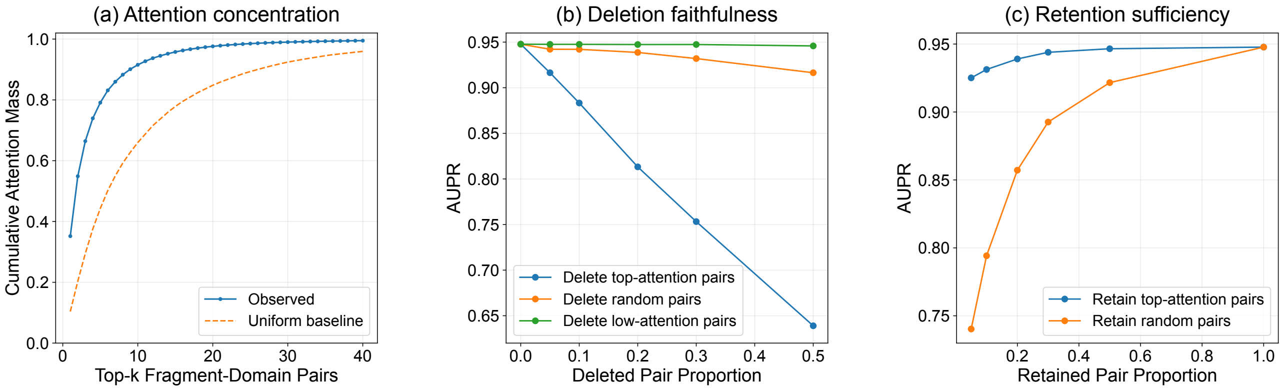


Figure 1: Attention concentration, deletion faithfulness, and retention sufficiency of fragment–domain interactions learned by DQHTFI. (a) Cumulative attention mass of the top-$k$ fragment–domain pairs compared with a uniform-attention baseline. (b) AUPR after deleting high-attention, random, or low-attention pairs. (c) AUPR after retaining high-attention or random pairs.

**MolTrans** (Huang et al. 2021). MolTrans is an end-to-end architecture that applies a substructure partitioning strategy. It decomposes drugs and proteins into frequently occurring subsequences and employs Transformer-based encoders to model contextual representations and interactions between local substructures.

**DrugBAN** (Bai et al. 2023). DrugBAN introduces a bilinear attention network to explicitly model fine-grained pairwise local interactions between drug substructures and protein features. It further incorporates conditional domain adversarial learning to improve generalization across different data distributions.

**MotifGT-DTI** (Tian et al. 2026a). MotifGT-DTI employs graph Transformers to encode drug molecular motifs and protein pocket subgraphs. It uses cross-attention to fuse protein sequence and structural features and subsequently applies bilinear attention to model local drug–protein interactions.

**MMDG-DTI** (Hua et al. 2025). MMDG-DTI is a multimodal feature fusion and domain generalization method. It employs pretrained language models to extract semantic features and a hybrid graph neural network to extract structural features. Domain adversarial training and contrastive learning are further incorporated to enhance generalization to unseen domains.

**DeepDTA** (Öztürk, Özgür, and Ozkirimli 2018). DeepDTA is a classical sequence-based DTA prediction model. It separately employs CNNs to extract features from drug SMILES strings and protein sequences and predicts drug–target binding affinity through fully connected layers.

**MGraphDTA** (Yang et al. 2022). MGraphDTA is based on a deep multiscale graph neural network. It represents drugs as molecular graphs and employs a multiscale GNN to extract local and global drug structural information, while using a CNN to extract protein sequence features.

**MFCLDTA** (Tian et al. 2026b). MFCLDTA is a multiscale feature contrastive-learning model for DTA prediction. It integrates molecular sequence features, molecular structural features, and affinity-graph information and employs contrastive learning to enhance multiscale representation alignment.

**DeepDTAGen** (Shah et al. 2025). DeepDTAGen is a multitask DTA framework. It employs a drug graph encoder and a protein Gated-CNN to extract drug–target features and simultaneously performs affinity prediction and target-aware drug generation within a shared feature space.

**MT-DiffGen** (Li et al. 2026). MT-DiffGen is a multitask diffusion model that jointly performs drug–target affinity prediction and target-aware molecular generation. It uses affinity information to guide the molecular generation process.

### E DTI Attention Analysis

To evaluate whether the learned fragment–domain attention reflects the predictive importance of local interactions, we conduct attention concentration, deletion, and retention experiments on BindingDB. Fragment–domain pairs are ranked according to their attention weights and are selectively removed or retained at the final interaction aggregation stage. The results are shown in Figure 1.

As shown in Figure 1(a), the attention distribution is highly concentrated. The top five fragment–domain pairs account for more than 70% of the total attention, while the cumulative attention of the top ten pairs approaches 90%, substantially exceeding the uniform-allocation baseline.

Figure 1(b) shows that deleting high-attention pairs causes a considerably larger performance decrease than deleting random or low-attention pairs. When 50% of the high-attention pairs are removed, AUPR decreases from approximately 0.95 to 0.64, whereas removing low-attention pairs has little effect. Conversely, as shown in Figure 1(c), retaining only approximately 5% of the high-attention pairs preserves an AUPR of approximately 0.92, while retaining the same proportion of random pairs produces substantially lower performance.

These results demonstrate that the attention weights are concentrated on a small number of informative fragment–domain interactions and are consistent with their contributions to DTI prediction.

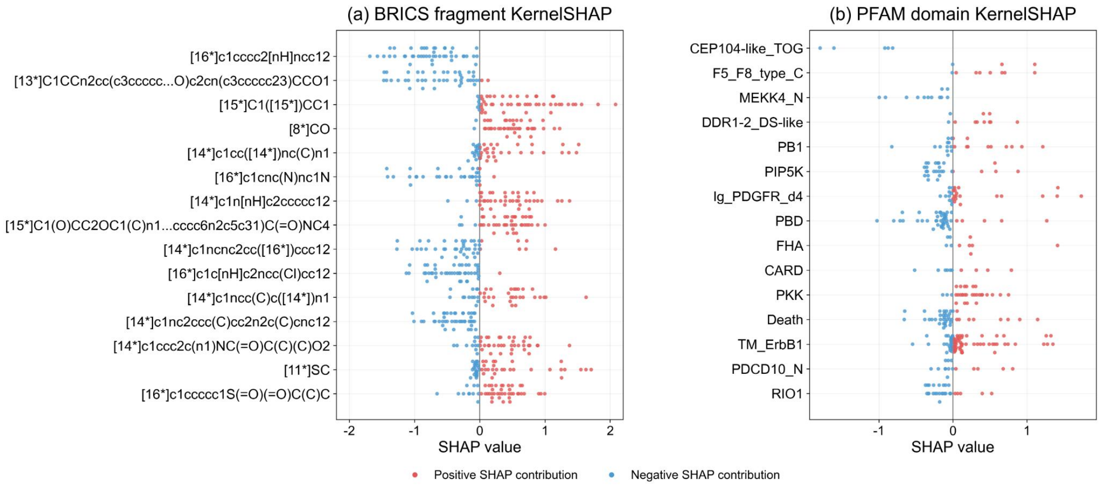


Figure 2: SHAP-based feature attribution for DQHTFI in DTA prediction. (a) SHAP values of BRICS drug fragments. (b) SHAP values of Pfam protein domains.

#### F DTA Feature Analysis

We further employ KernelSHAP (Lundberg and Lee 2017) on the Davis dataset to quantify the contributions of BRICS drug fragments and Pfam protein domains to DTA prediction. In Figure 2, positive and negative SHAP values indicate increases and decreases in the predicted affinity, respectively, while larger absolute values indicate stronger contributions.

For drug fragments, `[15*]C1([15*])CC1`, `[8*]CO`, and `[11*]SC` mainly exhibit positive contributions, whereas `[16*]c1cnc(N)nc1N` is primarily associated with negative contributions. Some fragments show both positive and negative SHAP values, indicating that their effects depend on the corresponding target and interaction context.

For protein domains, `CEP104-like_TOG`, `MEKK4_N`, and `RIO1` mainly exhibit negative contributions, whereas `F5_F8_type_C`, `PKK`, and `TM_ErbB1` show positive contributions in some samples. Domains such as `PB1`, `PBD`, and `Death` contribute in both directions, suggesting drug-dependent effects.

Overall, the attribution results indicate that DQHTFI dynamically integrates drug-fragment and protein-domain information rather than relying on a single fixed feature, providing chemically and biologically meaningful explanations for DTA prediction.

#### References


Bai, P.; Miljković, F.; John, B.; and Lu, H. 2023. Interpretable Bilinear Attention Network with Domain Adaptation Improves Drug–Target Prediction. *Nature Machine Intelligence*, 5: 126–136.

Davis, M. I.; Hunt, J. P.; Herrgard, S.; Ciceri, P.; Wodicka, L. M.; Pallares, G.; Hocker, M.; Treiber, D. K.; and Zarrinkar, P. P. 2011. Comprehensive Analysis of Kinase Inhibitor Selectivity. *Nature Biotechnology*, 29(11): 1046–1051.

Hua, Y.; Feng, Z.; Song, X.; Wu, X.-J.; and Kittler, J. 2025. MMDG-DTI: Drug–Target Interaction Prediction via Multimodal Feature Fusion and Domain Generalization. *Pattern Recognition*, 157: 110887.

Huang, K.; Xiao, C.; Glass, L. M.; and Sun, J. 2021. MolTrans: Molecular Interaction Transformer for Drug–Target Interaction Prediction. *Bioinformatics*, 37: 830–836.

Li, S.; Wang, H.; Hu, T.; Zhuang, L.; Zhao, J.; Sheng, Y.; and Sun, Y. 2026. MT-DiffGen: Unifying Affinity Prediction and Target-Aware Molecule Generation with a Multi-Task Diffusion Model. *Knowledge-Based Systems*, 340: 115605.

Liu, H.; Sun, J.; Guan, J.; Zheng, J.; and Zhou, S. 2015. Improving Compound–Protein Interaction Prediction by Building Up Highly Credible Negative Samples. *Bioinformatics*, 31(12): i221–i229.

Liu, T.; Lin, Y.; Wen, X.; Jorissen, R. N.; and Gilson, M. K. 2007. BindingDB: A Web-Accessible Database of Experimentally Determined Protein–Ligand Binding Affinities. *Nucleic Acids Research*, 35(suppl_1): D198–D201.

Lundberg, S. M.; and Lee, S.-I. 2017. A Unified Approach to Interpreting Model Predictions. In *Advances in Neural Information Processing Systems*, volume 30, 4765–4774. Curran Associates, Inc.

Nguyen, T.; Le, H.; Quinn, T. P.; Nguyen, T.; Le, T. D.; and Venkatesh, S. 2021. GraphDTA: Predicting Drug–Target Binding Affinity with Graph Neural Networks. *Bioinformatics*, 37(8): 1140–1147.

Öztürk, H.; Özgür, A.; and Ozkirimli, E. 2018. DeepDTA: Deep Drug–Target Binding Affinity Prediction. *Bioinformatics*, 34: i821–i829.

Shah, P. M.; Zhu, H.; Lu, Z.; Wang, K.; Tang, J.; and Li, M. 2025. DeepDTAGen: A Multitask Deep Learning Framework for Drug-Target Affinity Prediction and Target-Aware Drugs Generation. *Nature Communications*, 16: 5021.

Tang, J.; Szwajda, A.; Shakyawar, S.; Xu, T.; Hintsanen, P.; Wennerberg, K.; and Aittokallio, T. 2014. Making Sense of Large-Scale Kinase Inhibitor Bioactivity Data Sets: A

Comparative and Integrative Analysis. *Journal of Chemical Information and Modeling*, 54(3): 735–743.

Tian, W.; Zeng, M.; Wang, J.; and Lu, C. 2026a. MotifGT-DTI: Pivotal Motif-Based Graph Transformer Model Improves Drug–Target Interaction Prediction. *IEEE Transactions on Neural Networks and Learning Systems*, 1–15.

Tian, Z.; Zhu, S.; Teng, Z.; Yan, X.; and Wang, T. 2026b. MFCLDTA: Multi-Scale Feature Contrastive Learning for Predicting Drug-Target Binding Affinity. *Expert Systems with Applications*, 306: 130918.

Yang, Z.; Zhong, W.; Zhao, L.; and Chen, C. Y.-C. 2022. MGraphDTA: Deep Multiscale Graph Neural Network for Explainable Drug–Target Binding Affinity Prediction. *Chemical Science*, 13: 816–833.

Zhao, Q.; Zhao, H.; Zheng, K.; and Wang, J. 2022. HyperAttentionDTI: Improving Drug–Protein Interaction Prediction by Sequence-Based Deep Learning with Attention Mechanism. *Bioinformatics*, 38: 655–662.

Zitnik, M.; Sosič, R.; Maheshwari, S.; and Leskovec, J. 2018. BioSNAP Datasets: Stanford Biomedical Network Dataset Collection. Stanford Biomedical Network Dataset Collection.
